# Balances: a new perspective for microbiome analysis

**DOI:** 10.1101/219386

**Authors:** J. Rivera-Pinto, J.J. Egozcue, V. Pawlowsky–Glahn, R. Paredes, M. Noguera-Julian, M. L. Calle

## Abstract

High-throughput sequencing technologies have revolutionized microbiome research by allowing the relative quantification of microbiome composition and function in different environments. One of the main goals in microbiome analysis is the identification of microbial species that are differentially abundant among groups of samples, or whose abundance is associated with a variable of interest. Most available methods for microbiome abundance testing perform univariate tests for each microbial species or taxa separately, ignoring the compositional nature of microbiome data.

We propose an alternative approach for microbiome abundance testing that consists on the identification of two groups of taxa whose relative abundance, or balance, is associated with the response variable of interest. This approach is appealing, since it has direct translation to the biological concept of ecological balance between species in an ecosystem. In this work, we present *selbal*, a greedy stepwise algorithm for balance selection. We illustrate the algorithm with 16s abundance data from an HIV-microbiome study and a Crohn-microbiome study.

**Importance:** A more meaningful approach for microbiome abundance testing is presented. Instead of testing each taxon separately we propose to explore abundance balances among groups of taxa. This approach acknowledges the compositional nature of microbiome data.

## INTRODUCTION

Human microbiome research, focused on understanding the role in health and disease of microbes living in the human body, has experienced significant growth in the last few years. High-throughput sequencing technologies have revolutionized this field by allowing the quantification of microbiome composition and function in different environments. Large scale projects, like the Human Microbiome Project (1),(2) or MetaHIT (Metagenomics of the Human Intestinal Tract), have established standardized protocols for creating, processing and interpreting metagenomic data (3). However, the analysis of microbiome data for differential abundance or association with sample metadata is still challenging.

Typically, after DNA sequencing, bioinformatics preprocessing and quality control of the sequences, an abundance table with the number of sequences (*reads*) per sample for different microbial species (taxa) is obtained. Total number of sequences for each sample is highly variable, and depends on laboratory sample preparation. Indeed, raw abundances and the total number of reads per sample are non-informative since they depend on physical and technical mechanisms when sequencing the DNA. In order to mitigate the problem of different sampling depth, microbiome data are often normalized previous to differential abundance testing (4),(5).

Working with proportions, that is, relative abundances instead of raw abundances, does not solve the problem since there is a dependence structure in the data that may lead to misleading results such as spurious correlations or incoherent distances (6),(7). Rarefaction, which consists of random sampling of the same number of sequences for each sample, is similar to working with relative abundances. Though rarefaction might be convenient for richness and diversity analyses and avoids the problem of different sample depth, it supposes a loss of information and the increase of Type I error for differential abundance analyses (4). Other normalization alternatives, developed for RNA-Seq, are also applied in microbiome analysis for dealing with the problem of different number of counts per sample through variance stabilizing transformations (5). However, these RNA-Seq proposals also present problems with the false discovery rate when library sizes are very different among samples (8).

An alternative approach to rarefaction and normalization methods for microbiome analysis is to acknowledge the compositional nature of microbiome data and to use the mathematical theory available for compositional data (CoDa). Compositional data is defined as a vector of strictly positive real numbers carrying relative information. Relative information refers to the fact that the information of interest is contained in the ratios between the components of the composition and the numerical value of each component by itself is irrelevant (9).

As mentioned before, raw microbiome abundances are by itself non-informative since they depend on technical artifacts such as sequencing depth. Thus, microbiome data fits the definition of compositional data except for the fact that microbiome abundance tables contain many zeros. Assuming that observed zeros are rounded zeros, meaning that they correspond to values below the detection limit, they can be replaced by a positive value or pseudo count (10) so that CoDa analysis in terms of relative abundances between groups of microorganisms can be applied.

Several recent works acknowledge the compositional nature of microbiome abundance data and propose their analysis accordingly (11,12). Most of these approaches consider the *centered log-ratio transformation* (clr) and perform relative abundance testing for each clr transformed component, which is given by the logarithm of the component divided by the geometric mean of all the components in the sample. This allows the identification of clr transformed components that are associated with a specific characteristic of interest. However, the interpretation of such association is not straightforward because the clr transformation involves the abundances of all the taxa in the sample.

Instead, we propose to perform microbiome relative abundance testing by identifying two groups of taxa whose relative abundance is associated with the phenotype of interest. For this we use the notion of balance between two groups of components of a composition, which is a central concept in CoDa analysis.

Mathematically, a balance is defined as follows. Let *X* = (*X*_1_,*X*_2_, …,*X_k_*) be a composition of the number of counts for k different microbial species or taxa. Given two disjoint subsets of components in *X*, denoted by *X*_+_ and *X*_−_, indexed by *I*_+_ and *I*_−_, and composed by *k*_+_ and *k*_−_ taxa, respectively, the balance between *X*_+_ and *X*_−_ is defined as:

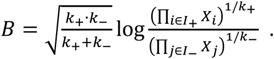

Expanding the logarithm, the balance is proportional to

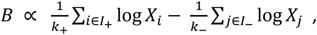

which is a more familiar expression corresponding to the difference in means of the log-transformed abundances between the two groups.

Balances are in compositional data analysis a key element in the construction of new coordinates through the so called *isometric log-ratio transformation* (ilr) (13) .

The concept of balance, as proposed in the compositional data theory, provides a new and interesting perspective for microbiome data analysis, since this mathematical concept is related to the biological concept of ecological balance in ecosystems.

Recently, some authors have proposed the use of CoDa approaches for microbiome analysis with different objectives such as the differential abundance between groups (14), differentiation of niches (15), or the inclusion of phylogenetic associations between the components included in the study (16).

In this work, we propose an algorithm for the identification of balances between groups of taxa that are associated with a dependent component of interest. This approach provides a new perspective to differential abundance and microbiome association studies. Starting with the balance composed by only two taxa that is most associated with the response, the algorithm performs a forward selection process and, at each step, a new taxon is added to the existing balance so that the specified association criterion is maximized. The algorithm stops when none of the possible additions improves the current association.

The paper is organized as follows. In the Results and Discussion section, the proposed algorithm is applied to an HIV-microbiome study and to a Crohn’s disease-microbiome study. Then these results are analyzed and both the advantages and technical issues of the algorithm when applied to microbiome data sets are discussed. Finally, in Material and Methods we present a detailed explanation of the algorithm.

## RESULTS

We illustrate the proposed methodology with a dataset from a cross – sectional HIV – microbiome study conducted in Barcelona (Spain) including both HIV – infected subjects and HIV – negative controls (17). Microbiome information is derived from MiSeq™ 16SrRNA sequence and bioinformatically processed with Mothur. After applying abundance filters and agglomerating taxa to genus level, microbiome abundance is summarized in a matrix of raw abundances for 155 samples and 60 different genera (Bioproject accession number: PRJNA307231, SRA accession number: SRP068240). Below, we present the results for two different analyses, the association of microbiome abundance with HIV status and with the inflammation parameter, sCD14. In the first case, the component of interest is dichotomous while in the second case it is continuous.

We also present the results of a Crohn’s disease study (18). Only patients with Crohn’s disease (*n* = 662) and those without any symptom (*n* = 313) were analyzed. The information was obtained from MiSeq™ 16SrRNA sequence, agglomerated to the genus level, resulting in a matrix with information of 48 genus for 975 samples. In this case, the goal is to identify groups of taxa whose abundance balance is associated with Crohn’s disease.

### Microbiome and HIV status

The main goal of this analysis is to find a microbiome balance associated with HIV-status, that is, a microbiome balance that is able to discriminate between HIV-positive and HIV-negative individuals. As exposed in Noguera–Julian et al. (17), the HIV risk factor MSM (Men who have Sex with Men) vs non-MSM should be considered as a possible confounder in any HIV - microbiome study. The proposed algorithm implements a regression model which allows adjustment for other variables. Thus, we applied the algorithm to *Y*=HIV-status and *X*=microbiome abundance at genus level, adjusted by *Z*=MSM factor.

According to the cross-validation (cv) procedure implemented with function *selbal.cv*, the optimal number of components to be included in the balance is 2 (Figure 1). The balance we identified as the most associated with HIV-status is given by *X*_+_, a taxon of the family *Erysipelotrichaceae* and unknown genus and *X*_−_, a taxon of the family *Ruminococcaceae* and unknown genus (Figure 2). HIV-positive status is associated with higher balance scores, that is, larger relative abundances of *Erysipelotrichaceae* with respect to *Ruminococcaceae.* The discrimination accuracy of this balance is moderate, with an AUC of 0.786 on the whole sample and a cross-validation AUC of 0.674. As can be observed in the boxplot in Figure 2, HIV-negative individuals are associated with lower balance values, most of them negative, while HIV-positive individuals have heterogeneous balance values. Figure 3 shows the result of the cross – validation procedure. The balance identified with the whole dataset is the most frequently identified in the cross-validation procedure, appearing 44% of the times, an indicator of robustness for the proposed global balance.

**Figure 1:**
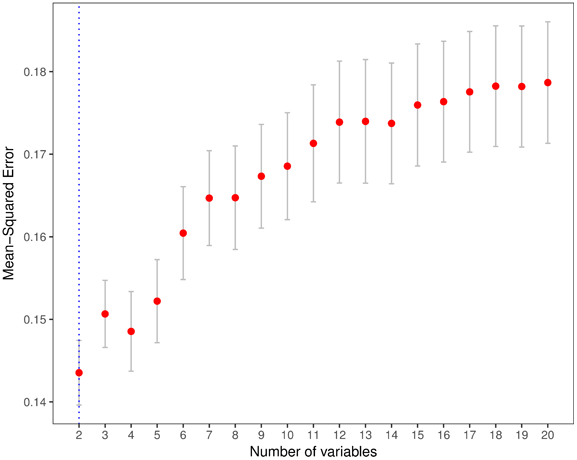
Mean squared error (MSE) as a function of the number of components included in the balance. The optimal number of components is highlighted with a vertical dashed line.

**Figure 2:**
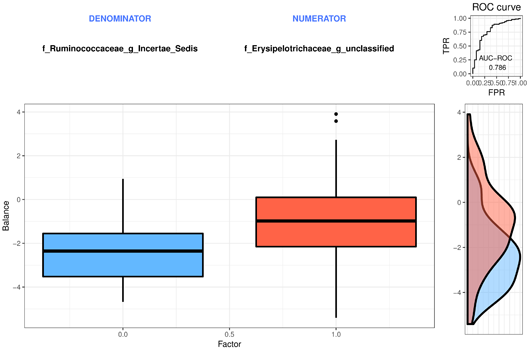
The components defining the selected balance are specified on top of the boxplot that represents the distribution of the balance score for each of the groups. The right part of the figure contains the ROC–curve with its AUC value (0.786) and the density curve for each group.

**Figure 3:**
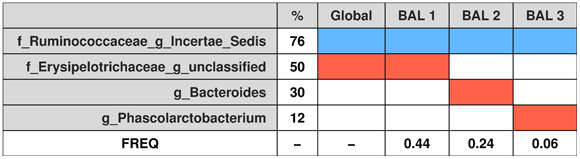
Cross–validation (cv) results: first column contains the names of the taxa appearing in the most frequently selected balances in the cv procedure, the second column provides the frequency of selection (in percentage), the third column corresponds to the global balance, that is, the balance obtained using all the samples. Columns 4 to 6 represent the most frequent balances identified in the cv procedure. Colored rectangles indicate if the component is in the numerator of the balance (*red*), in the denominator (*blue*) or not included (*white*). The last row provides the proportion of times the balance has been selected as optimal in the cv procedure.

### Microbiome and sCD14 inflammation parameter

Acute and chronic inflammations typically occur after HIV infection. Even patients under antiretroviral medications and undetectable viral load present chronic inflammation, which may cause tissue damage and is associated with many chronic diseases. In this context, there is a great interest in defining possible interventions involving modifications of the gut bacterial environment, which may reduce inflammation in HIV patients. This requires a good understanding of the association between gut microbial composition and several inflammation parameters. In this case, we focus on an immune–marker related to the chronic inflammation: the levels of soluble CD14 (sCD14), which was measured for a subset of samples (*n* = 151). The optimal number of components to be included in the model is four, according to the cv-MSE (Figure 4). The balance that is identified as the most associated with sCD14 is composed by two taxa in the numerator, *X*_+_ = {*g_Subdoligranulum, f_Lachnospiraceae_g_unclassified*} and two in the denominator *X*_−_ = {*f_Lachnospiraceae_g_Intertae_Sedis, g_Collinsella*}. The association is moderate, with R = 0. 53. Figure 5 provides a scatter plot of the balance values and sCD14 values, indicating that higher balance scores are associated with higher sCD14 values. The robustness of the selected balance can be evaluated through the results of the cv-procedure (Figure 6). We see that the proposed global balance is also the one that has been more frequently selected in the cv, 34% of the times. The four taxa defining the global balance correspond to the top 4 most frequently selected in the cross - validation. These results emphasize the robustness of the selected global balance.

**Figure 4:**
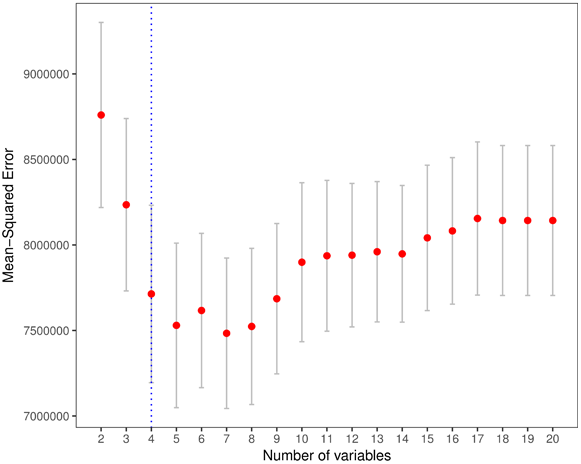
Mean squared error (MSE) as a function of the number of components included in the balance. The optimal number of components is highlighted with a vertical dashed line.

**Figure 5.**
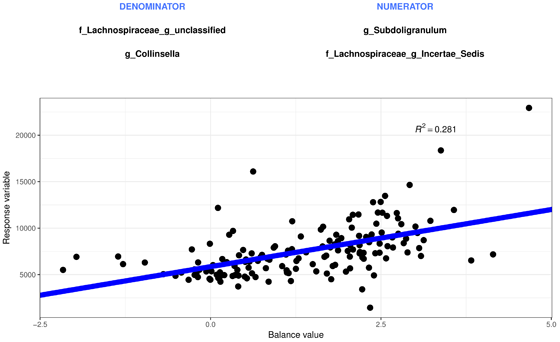
Representation of the balance obtained (*X* axis) for the sCD14 immune-marker values (*Y* axis), the bacteria groups composing it (top of the figure) and the corresponding regression line (*blue*).

**Figure 6:**
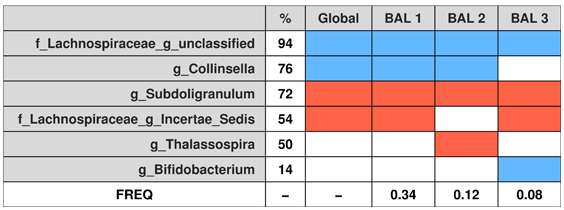
Cross – validation (cv) results: first column contains the names of the taxa included in the most frequently selected balances in the cv procedure, the second column provides the frequency of selection (in percentage), the third column corresponds to the global balance, that is, the balance obtained using the whole sample. Columns 4 to 6 represent the most frequent balances identified in the cv procedure. Colored rectangles indicate if the component is in the numerator of the balance (*red*), in the denominator (*blue*) or not included (*white*). The last row provides the proportion of times the balance has been the selected in the cv procedure.

### Crohn’s disease

Crohn’s disease is an inflammatory bowel disease (IBD) linked to microbial alterations in the gut (18),(19). We ran *selbal.cv* algorithm with the goal of identifying groups of taxa whose abundance balance can discriminate between individuals with Crohn’s disease from those without the disease.

The optimal number of components in the balance is twelve according to the MSE criterion (Figure 7). The groups defining the balance are *X*_+_= {*g_Roseburia, o_Clostridiales_g_, g_Bacteroides, f_Peptostreptococcaceae_g_*} and *X*_−_ = {*g_Dialister, g_Dorea, o_Lactobacillales_g_, g_Eggerthella, g_Aggregatibacter, g_Adlercreutzia, g_Streptococcus, g_Oscillospira*}. Cases with Crohn’s disease have lower balance scores than controls (Figure 8) which means lower relative abundances of *X*_+_ with respect to *X*_−_. The discrimination value of the identified balance is important, with an AUC = 0.838 and a cv-AUC = 0.819.

**Figure 7:**
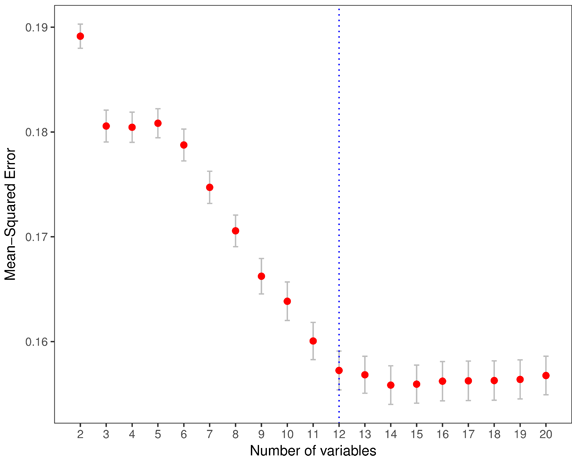
Mean squared error (MSE) as a function of the number of components included in the balance. The optimal number of components is highlighted with a vertical dashed line.

**Figure 8:**
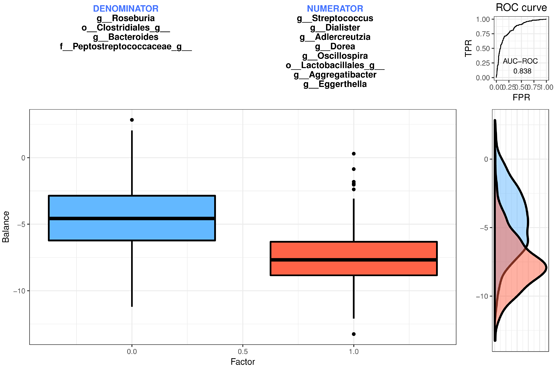
The components defining the selected balance are specified on top of the boxplot which represents the distribution of the balance score for each of the groups. The right part of the figure contains the ROC – curve with its AUC value (0.838) and the density curve for each group.

The identified global balance is very robust as the results of the cv reveal (Figure 9). The global balance obtained with the whole dataset is also the most frequently identified balance in the cv-procedure, namely 36% of the times. Moreover, the components defining the global balance are also the ones more frequently selected in the cv procedure. The balance identifies *Bacteroides* and *Clostridiales* as part of the denominator of the balance, which have also been identified previously as less abundant in Crohn’s disease individuals than in controls (18).

**Figure 9:**
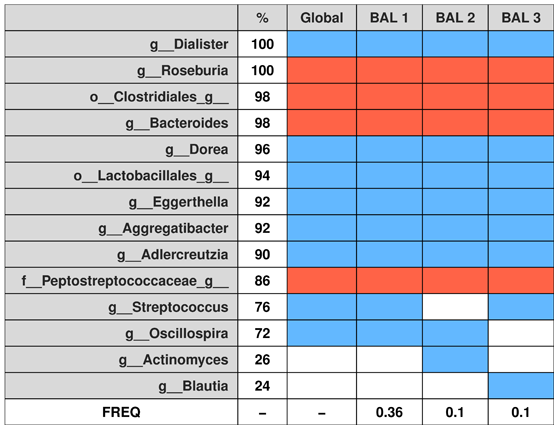
Cross – validation (cv) results: first column contains the names of the taxa appearing in the most frequently selected balances in the cv procedure, the second column provides the frequency of selection (in percentage), the third column corresponds to the global balance, that is, the balance obtained using the whole sample. Columns 4 to 6 represent the most frequent balances identified in the cv procedure. Colored rectangles indicate if the component is in the numerator of the balance (*red*), in the denominator (*blue*) or not included (*white*). The last row provides the proportion of times the balance has been the selected in the cv procedure.

## DISCUSSION

The identification of individual microbial species, or taxa, that are differentially abundant among groups of samples is challenging because the change in relative abundance of one taxon affects the relative abundances of the other taxa. As an alternative, we propose the analysis of relative abundances among groups of taxa instead of analyzing each taxon separately. In this work, we present *selbal*, a greedy stepwise algorithm for balance selection that takes into account the compositional nature of microbiome abundance data. The algorithm identifies two groups of taxa whose relative abundance, or balance, is associated with the response variable of interest.

*selbal* overcomes the problem of differences in sample size that is usually treated with different methods based on count-normalization, rarefaction or transformation into proportions. The only way in which data is altered in *selbal* is at the zero imputation stage required because of the use of logarithms and ratios in the definition of balances. This replacement of zeros by positive numbers is performed under the assumption that observed zeros are rounded zeros, that is, all taxa are present in all the samples but some of them are not detected because of low abundance and insufficient sample depth. However, it is not clear how the imputation method and the presence of structural zeros (absence of the taxa in the sample) may influence the results. Future research will be focused on the treatment of zeros with the aim of more precisely evaluating if zeros are rounded or structural and on selecting the best replacement method.

Due to the computational cost, *selbal* does not explore the whole balance space and the method for selecting the optimal balance is suboptimal and may be improved. Thus, exploring for alternative approaches in the search of the optimal balance is another topic of future research.

In order to improve classification or prediction accuracy of the variable of interest a prediction model with several balances can be obtained by applying *selbal* algorithm sequentially. This sequential search of balances can be performed similarly to partial least squares approach: when the first balance B1 is identified, all variables are deflated by the first balance, that is, each variables is adjusted for the first balance, by regressing the variable on B1 and taking residuals. Then, the second balance is searched on the new orthogonalized data.

Endorsed by the compositional treatment of microbiome abundance data, *selbal* can also be useful for comparing different microbial studies. Since balances are based on relative abundances among groups of taxa, this relative information is likely to remove the noise and biases of each particular study.

## MATERIALS AND METHODS

Let *X* = (*X*_1_,*X*_2_,…,*X_k_*) be a composition, that is, a vector of strictly positive real numbers. Given two disjoint subsets of components in *X*, denoted by *X*_+_ and *X*_−_, indexed by *I*_+_ and *I*_−_, composed by *k*_+_ and *k*_−_ components respectively, the balance between *X*_+_ and *X*_−_is defined as:

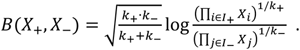

Expanding the logarithm, the balance is proportional to

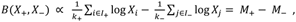

which corresponds to the difference of the arithmetic means of the log-transformed initial components in the two groups that we denote by *M*_+_ and *M*_−_, respectively. This second expression is preferable from a computational point of view and is the one implemented in the proposed algorithm.

Given *Y*, a response variable, which can be either numeric or dichotomous, a composition *X* = (*X*_1_,*X*_2_,…,*X_k_*) and additional covariates *Z* = (*Z*_1_,*Z*_2_,…,*Z_r_*), the goal of the algorithm is to determine the sub-compositions of *X*, *X*_+_ and *X*_−_, indexed by *I*_+_ and *I*_…_, respectively, so that the balance *B* between *X*_+_ and *X*_−_ is highly associated with *Y* after adjustment for covariates *Z*. Depending on the nature of the dependent variable, the association can be defined in several ways.

For a continuous variable *Y*, the optimization criterion is defined as maximization of the coefficient of determination of the linear regression model:

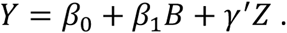

For a dichotomous variable *Y*, we fit the logistic regression model

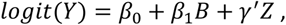

and, in this case, we consider three possible optimization criteria: the area under the ROC curve (default option), the maximization of the explained variance (20) or the discrimination coefficient (21).

The main function of the proposed algorithm to detect the most associated balance is called *selbal* and follows these steps:

**STEP 0:** *Zero replacement*

The initial matrix of counts in a microbiome study, denoted by 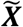, typically contains zeros. In order to apply the mathematical theory of compositional data, the observed zeros are assumed to be non-structural zeros but a consequence of under detection limit. They are replaced by a positive value using a Bayesian-Multiplicative replacement (10) of count zeros as implemented in function *cmultRep()* of the R package *zCompositions* (22). It is important to remark that this transformation keeps the information contained in the ratios between non-zero components. The resulting matrix without zeros is denoted by ***X*** and coincides with 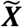 only if the latter has no null value.

**STEP 1:** *Optimal balance between two components*

The algorithm evaluates exhaustively the optimization criterion for all possible balances composed by only two components; that is, all the balances of the form:

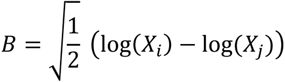

for *i,j* ∈ {1,…, *k*} *i* ≠ *j*. We denote by *B*^(1)^ the optimal two-component balance in terms of maximization of the association value.

For each pair of components (*X_i_*,*X_j_*) there are two options when defining a balance:

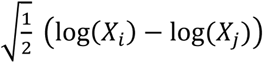

and

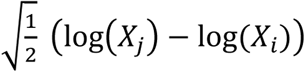

differenced only by their sign. For dichotomous variables, they will provide the same AUC value; nevertheless *selbal* returns the balance whose coefficient in the regression model is positive.

**STEP *s*:** *Optimal balance adding a new* component

For *s* > 1 and until the stop criterion is fulfilled, let *B*^(*s*−1)^ be

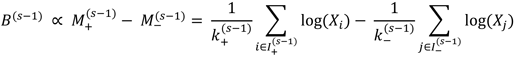

where 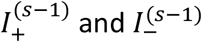 are two disjoint subsets of indices in {1, …, *k*}, with 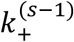 and 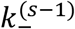 elements, respectively.

For each index 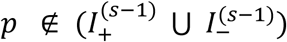, the algorithm evaluates the optimization criterion of the balance that is obtained by adding log(*X_p_*) to *B*^(*s*−1)^ including *p* either in 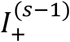 or in 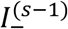. That is, the algorithm evaluates the optimization criterion for both, *B*^(*s*+)^ and *B*^(*s*−)^, defined as:

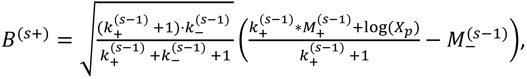

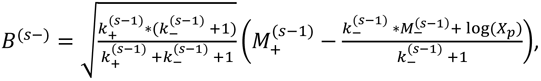

and selects as *B*^(*s*)^ the one that maximizes the optimization criterion. If *B*^(*s*)^ = *B*^(*s*+)^ the sets of new indices are defined as, 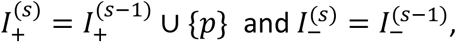 and similarly for *B*^(*s*)^ = *B*^(*s*−)^.

**STOP criterion.** *selbal* function has two parameters to decide the stopping criterion:

- *th.imp*, threshold improvement (default 0). The algorithm stops the iteration process when the improvement in association is lower than the specified threshold improvement.
- *maxV*, maximum number of components. The algorithm stops when the specified maximum number of components has been included in the balance.

### Cross-validation: *selbal.cv*

We perform a cross-validation procedure with two goals: (1) to identify the optimal number of components to be included in the balance and (2) to explore the robustness of the global balance identified with the whole dataset.

The cv procedure is implemented in the *selbal.cv* function.

For each cv process, the dataset is divided into K folds (default value, K = 5). K-1 folds are used to obtain the balance (with *th.imp* = 0 as the stop rule) and the remaining fold is used to test the result. The process is repeated M times (default value, M = 10)

### Optimal number of components

For each combination of K and M we perform the *selbal* function on the training dataset and find the optimal balance with *c* components, *c* ∈ {2,…,*C*} (default value, C = 20) and evaluate the mean squared error (MSE) of the model on the test dataset. For each *c* we obtain 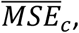 the mean MSE of the different models with *c* components and the corresponding standard error. The optimal number of components is defined with the 1se rule, as the minimum number of components whose mean MSE is below the minimum 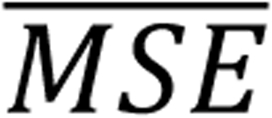 plus its standard error.

For dichotomous components, the MSE is computed in the same way codifying the two groups as 0 and 1.

### Robustness of the result

Once the optimal number of components *k_opt_* has been chosen, all the balances obtained in the cv procedure are reduced to *k_opt_* components. Then, a frequency table is built both for balances and for individual components. This information, available in the output of *selbal.cv*, is summarized in a table as those shown in Figure 3, Figure 6 and Figure 9.

The cv process also provides the association or discrimination value for each balance in the cv which can be used as a more accurate measure of association or discrimination of the global model.

## BIBLIOGRAPHY

1. Arumugam M, Raes J, Pelletier E, Paslier D, Yamada T, Mende DR, et al. Enterotypes of the human gut microbiome. Nature [Internet]. 2011 May 12 [cited 2014 Jul 9]; 473(7346):174–80. Available from: http://eutils.ncbi.nlm.nih.gov/entrez/eutils/elink.fcgi?dbfrom=pubmed&id=21508958&retmode=ref&cmd=prlinks

2. Turnbaugh PJ, Ley RE, Hamady M, Fraser-Liggett CM, Knight R, Gordon JI. The human microbiome project. Nature [Internet]. 2007 Oct 18 [cited 2014 Jul 10]; 449(7164):804–10. Available from: http://www.pubmedcentral.nih.gov/articlerender.fcgi?artid=3709439&tool=pmcentrez&rendertype=abstract

3. Santiago A, Panda S, Mengels G, Martinez X, Azpiroz F, Dore J, et al. Processing faecal samples: a step forward for standards in microbial community analysis. BMC Microbiol [Internet]. 2014 Jan [cited 2015 Aug 8];14(1):112. Available from: http://www.biomedcentral.com/1471-2180/14/112

4. McMurdie PJ, Holmes S. Waste Not, Want Not: Why Rarefying Microbiome Data Is Inadmissible. PLoS Comput Biol. 2014;10(4).

5. Dillies MA, Rau A, Aubert J, Hennequet-Antier C, Jeanmougin M, Servant N, et al. A comprehensive evaluation of normalization methods for Illumina high-throughput RNA sequencing data analysis. Brief Bioinform. 2013;14(6):671–83.

6. Pearson K. Mathematical Contributions to the Theory of Evolution. --On a Form of Spurious Correlation Which May Arise When Indices Are Used in the Measurement of Organs. 1896;

7. Gloor GB, Wu JR, Pawlowsky-Glahn V, Egozcue JJ. It’s all relative: analyzing microbiome data as compositions. Ann Epidemiol [Internet]. 2016;26(5):322–9. Available from: http://dx.doi.org/10.10167j.annepidem.2016.03.003

8. Weiss S, Xu ZZ, Peddada S, Amir A, Bittinger K, Gonzalez A, et al. Normalization and microbial differential abundance strategies depend upon data characteristics. Microbiome [Internet]. 2017;5(1):27. Available from: http://microbiomejournal.biomedcentral.com/articles/10.1186/s40168-017-0237-y

9. Pawlowsky-Glahn V, Egozcue JJ, Tolosana-Delgado R. Modeling and Analysis of Compositional Data. 2015. 272 p.

10. Martín-Fernández JA, Hron K, Templ M, Filzmoser P, Palarea-Albaladejo J. Bayesian-multiplicative treatment of count zeros in compositional data sets. Stat Modelling. 2015;15(2):134–58.

11. Mandal S, Treuren W Van, White RA, Eggesbø M, Knight R, Peddada SD. Analysis of composition of microbiomes: a novel method for studying microbial composition. 2015;1:1–7.

12. Cao KL, Costello M, Lakis VA, Bartolo F. MixMC : A Multivariate Statistical Framework to Gain Insight into Microbial Communities. 2016;

13. Egozcue JJ, Pawlowsky-Glahn V, Mateu-Figueras G, Barceló-Vidal C. Isometric Logratio Transformations for Compositional Data Analysis. Math Geol. 2003;35(3):279–300.

14. Fernandes AD, Reid JN, Macklaim JM, McMurrough TA, Edgell DR, Gloor GB. Unifying the analysis of high-throughput sequencing datasets: characterizing RNA-seq, 16S rRNA gene sequencing and selective growth experiments by compositional data analysis. Microbiome [Internet]. 2014;2:15. Available from: http://www.pubmedcentral.nih.gov/articlerender.fcgi?artid=4030730&tool=pmcentrez&rendertype=abstract

15. Morton JT, Sanders J, Quinn RA, Mcdonald D, Gonzalez A, Vázquez-baeza Y, et al. crossm Differentiation. 2017;2(1):1–11.

16. Washburne AD, Silverman JD, Leff JW, Bennett DJ, Darcy JL, Mukherjee S, et al. Phylogenetic factorization of compositional data yields lineage-level associations in microbiome datasets. PeerJ [Internet]. 2017;5:e2969. Available from: https://peerj.com/articles/2969

17. Noguera-Julian M, Rocafort M, Guillén Y, Rivera J, Casadellà M, Nowak P, et al. Gut Microbiota Linked to Sexual Preference and HIV Infection. EBioMedicine [Internet]. 2016;5:135–46. Available from: http://dx.doi.org/10.1016Zj.ebiom.2016.01.032

18. Ren B, Schwager E, Knights D, Song SJ, Yassour M, Haberman Y, et al. The treatment-naïve microbiome in new-onset Crohn’ s disease. 2015;15(3):382–92.

19. Øyri SF, Muzes G, Sipos F. Dysbiotic gut microbiome: A key element of Crohn’syri SF, Muzes G, Sipos F. Dysbiotic gut microbiome: A key element of Crohn’s disease. Vol. 43, Comparative Immunology, Microbiology and Infectious Diseases. 2015. p. 36–49.

20. Mittlböck M, Schemper M. Explained variation for logistic regression. Stat Med. 1996;15(19):1987–97.

21. Tjur T. Coefficients of Determination in Logistic Regression Models—A New Proposal: The Coefficient of Discrimination. Am Stat. 2009;63(4):366–72.

22. Martín-Fernández J, Barcelό-Vidal C, Pawlowsky-Glahn V. Dealing with Zeros and Missing Values in Compositional Data Sets Using Nonparametric Imputation. Math Geol. 2003;35(3):253–78.

